# Generating single-sex litters: development of CRISPR-Cas9 genetic tools to produce all-male offspring

**DOI:** 10.1101/2020.09.07.285536

**Authors:** Charlotte Douglas, Valdone Maciulyte, Jasmin Zohren, Daniel M. Snell, Obah A. Ojarikre, Peter J.I. Ellis, James M.A. Turner

## Abstract

Animals are extremely useful genetic tools in science and global resources in agriculture. However, a single sex is often required in surplus, and current genetic methods for producing all-female or all-male litters are inefficient. Using the mouse as a model, we developed a synthetic, two-part bicomponent strategy for generating all-male litters. We achieved this using CRISPR-Cas9 genome editing technology to generate large stable knock-ins on the autosomes and X chromosome. The bicomponent system functions via the sex-specific co-inheritance of a Cas9 transgene and an sgRNA transgene targeting the essential *Topoisomerase 1* gene. This technology proved to be highly efficient in generating on-target mutations, resulting in embryonic lethality of the target sex. Our study is the first to successfully generate all-male mammalian litters using a CRISPR-Cas9 bicomponent system and provides great strides towards generating single-sex litters for laboratory or agricultural research.

## Introduction

Animals and animal products are utilised globally. However, a single sex is often required at surplus, at the expense of the non-required littermates. Although the “Reduction, Replacement and Refinement” (3Rs) guidelines^1^ promote efficient animal use, the production of the unrequired sex is generally unavoidable. The unrequired sex may also exhibit a severe phenotype, thereby precluding its use for other experimental purposes. The generation of male-only litters could be advantageous for research focused on male-specific biology such as testis development, or Y chromosome studies (reviewed in ^2^), or on behavioural or drug response studies^3–5^, where sex is considered an important biological variable. It could also be useful in agriculture, for example the beef-producing industry, which preferentially uses male rather than female cows, because males are faster growing. Moreover, sex-specific selection of all-male broods could potentially greatly contribute to invasive pest control methods. Hypothetically, releasing broods of sterile male-only litters could induce population collapse, as the gametes are non-functional or produce sub-fertile offspring^6^. In this strategy, male-dominated populations could be used for controlling the propagation of malarial parasites, or insects that destroy food crops. Hence, a genetic method to produce all-male litters would be extremely beneficial.

The production of single-sex litters relies on differences in male and female chromosome complement and gene expression. With rare exceptions^7–9^, in eutherian mammals, females have two X chromosomes (XX) whilst males have a single X and a single Y chromosome (XY). The heterogametic nature of the XY chromosomes in males ensures the unique inheritance of either sex chromosome in a sex-specific manner. Consequently, sex-chromosome linked transgenes will also be inherited sex-specifically. The advent of CRISPR-Cas9 genome editing, the process of RNA-guided Cas9 endonuclease-driven DNA double strand breaks, provided an unprecedented ease with which to generate mutations in a vast range of cells *in vitro* and *in vivo*^10^ and can be used to generate these sex-chromosome linked transgenes.

Co-inheritance of a sex-linked Cas9 transgene, and an autosomal single guide RNA (sgRNA) transgene targeting an essential gene, can induce mutations and non-viability in one sex. Previously, Zhang and colleagues engineered a female-lethal bicomponent system in silkworms by integrating a Cas9 transgene onto the female-specific W chromosome, and an sgRNA transgene targeting essential housekeeping gene *Bmtra2* onto an autosome^11^. The co-inheritance of the W-Cas9 and *Bmtra2*-targeting sgRNA transgene was therefore uniquely in daughters; inducing *Bmtra2* loss-of-function mutations, and production of all-male litters with 100% efficiency^11^. In mice, Yosef and colleagues published a variation of this technology by engineering a Y-linked sgRNA transgene targeting essential embryonic genes *Atp5b, Casp8* and *Cdc20*^12^. Co-inheritance of the Y-linked sgRNA with an autosomal Cas9^13^ resulted in male-specific CRISPR-Cas9 mutations in the target genes, causing male lethality and a female-bias offspring sex ratio skew. However some male pups were born, and some showed severe developmental defects^12^.

Currently, a mammalian bicomponent genetic system for producing all-male litters has not been generated. We therefore utilised CRISPR-Cas9 genome editing to create a synthetic female-lethal bicomponent system in mice. We generated an X-linked Cas9 transgene and a second autosome-linked sgRNA transgene, whereby the sgRNA targets essential housekeeping gene *Topoisomerase 1* (*Top1*). We show that co-inheritance of a paternal X-linked Cas9 transgene and *Top1* sgRNA transgene results in lethality specifically in daughters. Remarkably, the number of offspring derived from this bicomponent approach exceeds the expected 50% of that from control matings. This unexpected finding reveals a buffering system that operates during preimplantation development to maximise offspring number. Our study is the first report of producing all-male litters in the mouse by sex-specific CRISPR-Cas9 genetic methods.

## Results

### An *in vitro* CRISPR-Cas9 bicomponent system induces *Top1* mutations

For our lethal-guide experiments, we chose to target *Top1*, a highly conserved 21-exon gene with essential functions in DNA replication and repair^14^. *Top1* loss-of-function results in embryonic lethality at the 4-16 cell stage^15–17^. We designed sgRNAs targeting exon 15 (sgRNA1) and exon 16 (sgRNA2), which together encode the DNA-binding domain, and a third targeting exon two (sgRNA3), adjacent to the start codon, thereby hypothetically disrupting the *Top1* reading frame early within the coding sequence (Fig 1A,B). Each sgRNA was inserted into a plasmid vector (annotated “pLethal”) driven by a human U6 (hU6) promoter. pLethal also encoded a pCbh promoter-driven mCherry reporter, which acted as a proxy for sgRNA expression.

**Fig 1.**
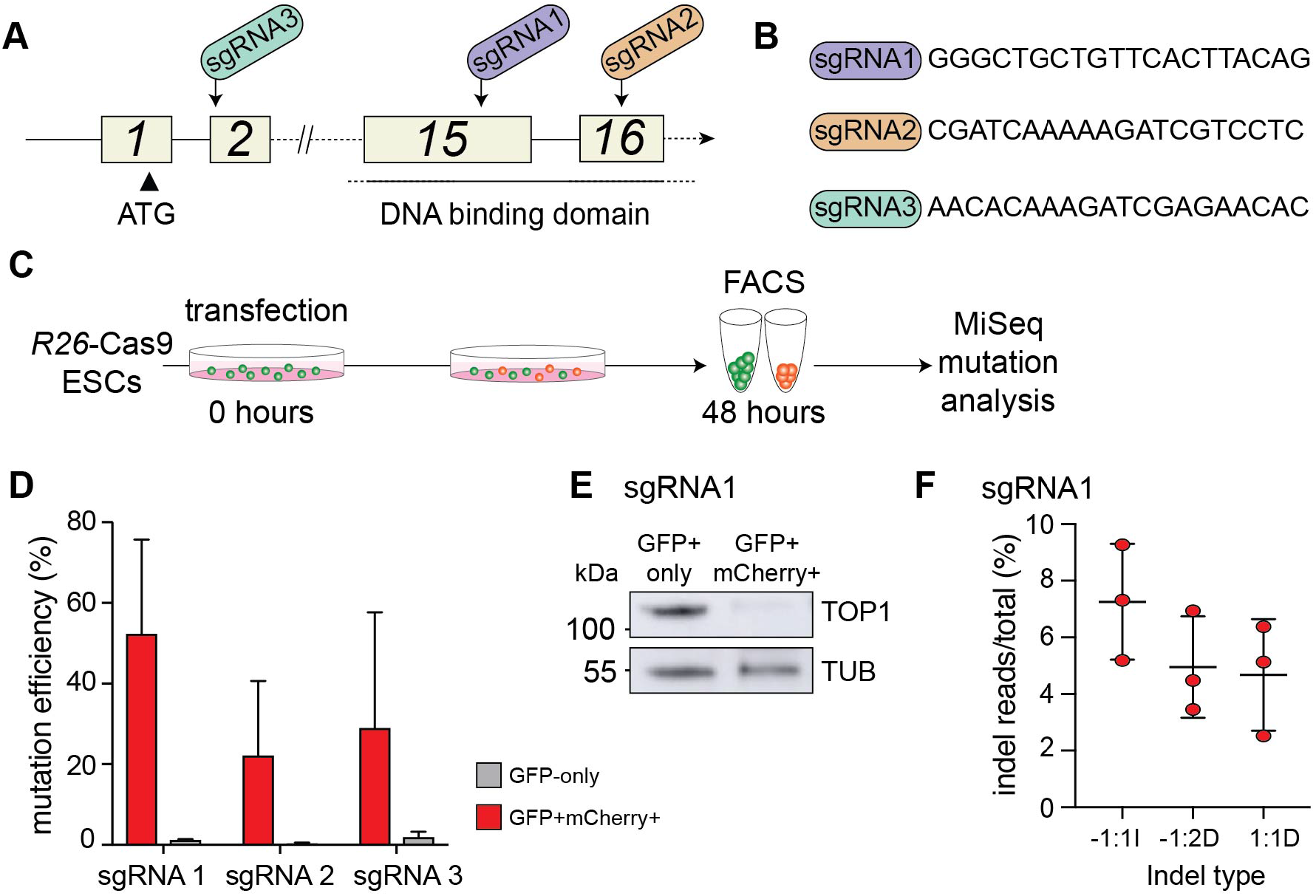
sgRNA1 Targeting *Top1* Exon 15 Induces the Greatest Mutation Efficiency. (A) Schematic of *Top1*, with single guide RNA (sgRNA) targeting essential exons. (B) Sequence of each 20mer sgRNA. (C) Schematic for screening sgRNAs in *R26*-Cas9 mESCs. (D) Quantification of mutation efficiency. Error bars: s.d. (n=3). (E) Western blot of GFP-only and double-positive mESCs after transfection with sgRNA1. Expected size: 110 kDa (TOP1) and 50 kDa (TUBULIN). (F) Occurrence of mutation types.

To screen individual sgRNAs in an *in vitro* bicomponent system, we derived constitutive pCAG promoter driven-Cas9 expressing XY mouse embryonic stem cell (mESC) lines from the *Gt(ROSA)26-Cas9* (*R26-Cas9*) transgenic mouse^13^. The Cas9 transgene is linked to an eGFP reporter via a T2A sequence. eGFP expression was confirmed by quantitative PCR (qPCR; Supp Fig 1A), and used as a visual proxy for Cas9 expression.

To assess whether the CRISPR-Cas9 bicomponent system could generate *Top1* mutations, *R26*-Cas9 mESCs were individually transfected with each sgRNA-containing pLethal plasmid and sorted 48 hours later by fluorescence-activated cell sorting (FACS; Fig 1C, Supp Fig 1B). mESCs transfected were eGFP and mCherry double-positive, while those not transfected were eGFP only. We evaluated the occurrence of *Top1* mutations in each population. sgRNA1 had the greatest mutation efficiency, with 52.24% of double-positive mESCs exhibiting *Top1* mutations (Fig 1D). sgRNA2 and sgRNA3 generated *Top1* mutations in 22.05% and 28.93% of double-positive mESCs, respectively (cf. 1.15%, 0.43% and 1.8% for sgRNA1,2 and 3 in eGFP-only cells, respectively; Fig 1D). sgRNA1 was carried forward for future experiments, since this guide exhibited the greatest mutagenic capacity. We confirmed by western blotting that in sgRNA1-transfected double-positive mESCs sgRNA1 TOP1 levels were reduced (Fig 1E). A single nucleotide insertion at the minus 1 position (−1:1I) downstream of the predicted Cas9 DSB site was the most dominant indel mutation, contributing on average 7.3% of all reads (Fig 1F). Moreover, this indel type aligned with the predicted mutational outcome for this sgRNA^18^. A −1:1I frame-shift mutation induces the occurrence of a premature stop codon; thereby fulfilling the requirement for a loss-of-function mutation.

### Co-inheritance of autosomal sgRNA and Cas9 transgenes induces *Top1* mutations and embryonic lethality

sgRNA1 generated frame-shift mutations at *Top1* exon 15, causing loss of TOP1. We therefore generated a transgenic mouse line, hereafter termed sgRNA*^Top1^*, that carried sgRNA1 together with the mCherry reporter cassette (Fig 2A). The sgRNA-mCherry transgene was integrated into an intergenic *Hipp11* (*H11*) locus on mouse chromosome 11 by ϕC31 integrase, which mediates efficient integration at attB and attP phage attachment sites^19,20^. For this knock-in, the sgRNA1 plasmid was edited to contain attB sequences flanking the U6-sgRNA1 and pCbh-mCherry, and was co-injected with integrase mRNA into the pronuclei of *H11*-attPx3 zygotes to generate founders (Fig 2B). mCherry expression was confirmed in sgRNA*^Top1^* mice by *in vivo* imaging of pups (Fig 2C), qPCR of adult tissues (Fig 2D) and fluorescence microscopy of pre-implantation embryos at embryonic day (E)3.5 (Fig 2E). The sgRNA*^Top1^* transgene did not induce embryonic lethality in isolation: breeding hemizygous sgRNA*^Top1^*/+ males with wildtype females (“ctrl”; Fig 2F) produced wildtype and sgRNA*^Top1^*/+ embryos in equal proportions (n=78; non-significant deviation from expected Mendelian ratio, p=0.65; Fig 2G).

**Fig 2.**
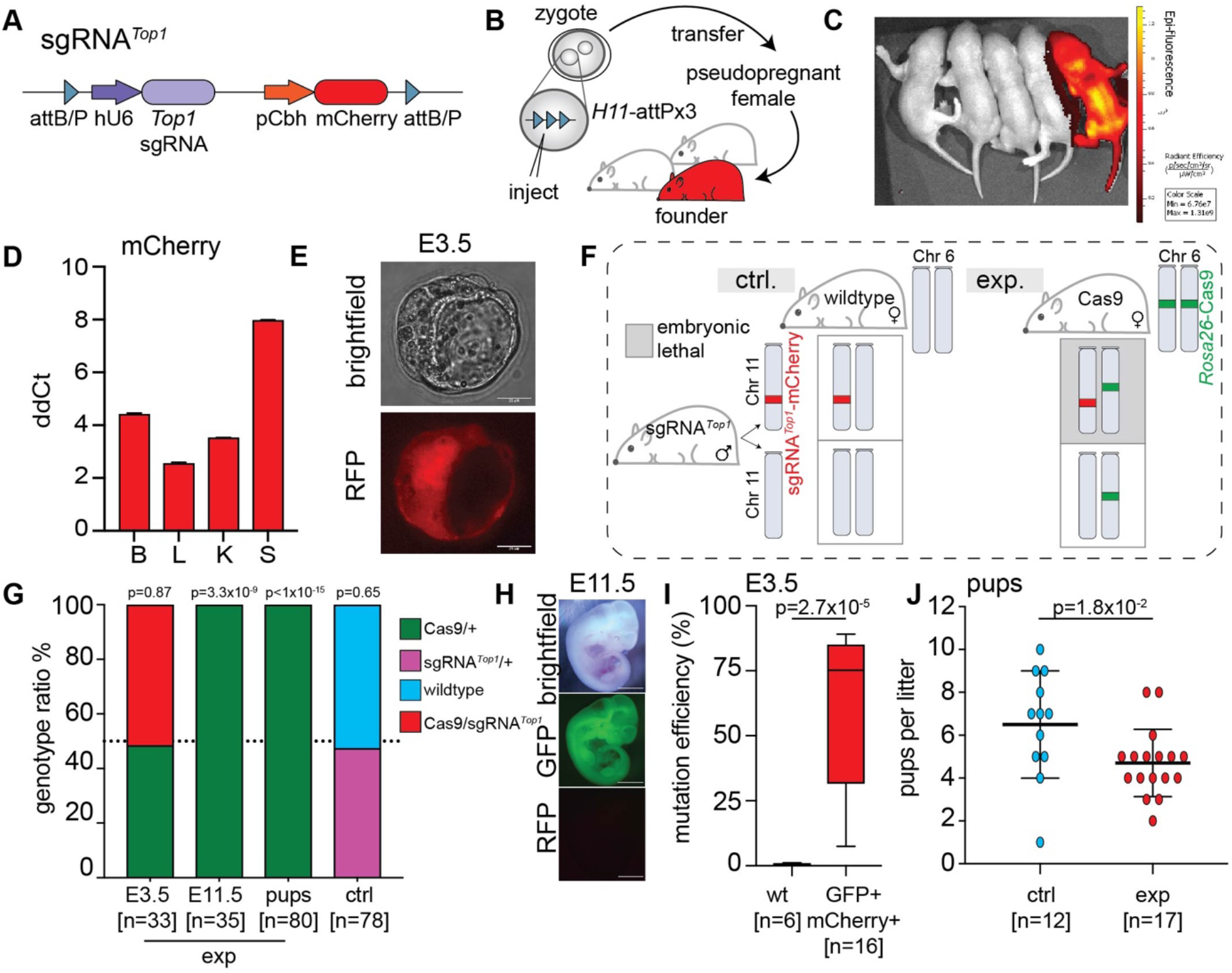
Characterising the sgRNA knock-in transgene. (A) Schematic of the *H11* sgRNA knock-in locus. (B) Schematic to generate the sgRNA*^Top1^* knock-in. (C) *In vivo* imaging for mCherry. (D) Quantitative PCR for mCherry expression normalised to *Gapdh* in a wildtype sample. B; brain, L; liver, K; kidney, S; spleen. Error bars: s.d. (n=3). (E) Fluorescence imaging of an sgRNA*^Top1^* E3.5 embryo. Scale bar; 25 μM. (F) Schematic of the mating strategies. ctrl; control, exp; experimental, Cas9; *R26*-Cas9. (G) Offspring genotypes during development from control or experimental matings. Number of offspring in brackets. P value: Chi-squared test, assuming 1:1 ratio of ‘expected’ offspring if co-inheritance of CRISPR-Cas9 alleles was not lethal. (H) Fluorescence imaging of E11.5 Cas9/+ embryo. Scale bar: 1mm. (I) Quantification of mutation efficiency. Error bars: range. Number of samples in brackets. P value: Mann-Whitney test. J) Litter size. Number of litters in brackets. P value: Mann-Whitney test.

To assess transgenic *Top1* sgRNA functionality *in vivo*, we bred hemizygous sgRNA*^Top1^*/+ males with homozygous *R26-*Cas9 females (“exp”; Fig 2F) and genotyped resulting embryos at multiple developmental stages (Fig 2G). In E3.5 blastocysts, the ratio of Cas9/ sgRNA*^Top1^* embryos to Cas9/+ was 1:1 (non-significant deviation from expected Mendelian ratio; p=0.87; Fig 2G). However, post-implantation, at E11.5, 100% of embryos were Cas9/+ (n=35; significant deviation from expected Mendelian ratio; p=3.3×10^−9^; Fig 2G,H). Later, at birth, all embryos were Cas9/+ (n=80; significant deviation from expected Mendelian ratio p<1×10^−15^; Fig 2G). In E3.5 eGFP/mCherry double-positive E3.5 embryos, the average mutation efficiency was 59.32% (n=16, cf. 0.65% in wildtype, p=2.7×10^−5^; Fig 2I). Moreover, all double-positive embryos exhibited the −1:1I mutation. Therefore, co-inheritance of a Cas9 and sgRNA transgene induced *Top1* mutations in pre-implantation embryos, and embryonic lethality with 100% efficiency prior to E11.5.

Due to the embryonic-lethal effect, we expected that the litter size in experimental matings would be 50% of that from control matings. Surprisingly however, this was not the case. Although the litter size in experimental matings was significantly reduced (4.7 versus 6.5; p=1.8×10^−2^), the mean litter size was 72% rather than 50% of controls (Fig 2J). This unexpected finding reveals a compensation mechanism operating *in utero* to maximise embryo number.

### Co-inheritance of an X-linked Cas9 transgene and autosome-encoded sgRNA*^Top1^* causes female lethality

To generate all-male litters using the CRISPR-Cas9 bicomponent system, we engineered an X-linked Cas9 transgenic mouse line containing a 3X FLAG-tagged Cas9 and eGFP reporter, linked via a T2A sequence and driven by a constitutive pCAG promoter (Fig 3A). The construct was targeted to the X-linked permissive *Hprt* locus, deletion of which has no effect on viability and fertility in mice^21–24^. Targeting in C57BL/6N mESCs generated a 173bp *Hprt* exon 2 deletion and a knock-in PCR product which we observed in 20% of mESC clones (n=9/48; Supp Fig 2A). We carried forward an X-Cas9 clone (“clone 5”) that by low-pass whole genome sequencing^25^ was confirmed to be euploid (Supp Fig 2B), gave rise to high-contribution chimeras from blastocyst injection, and germline transmitted.

**Fig 3.**
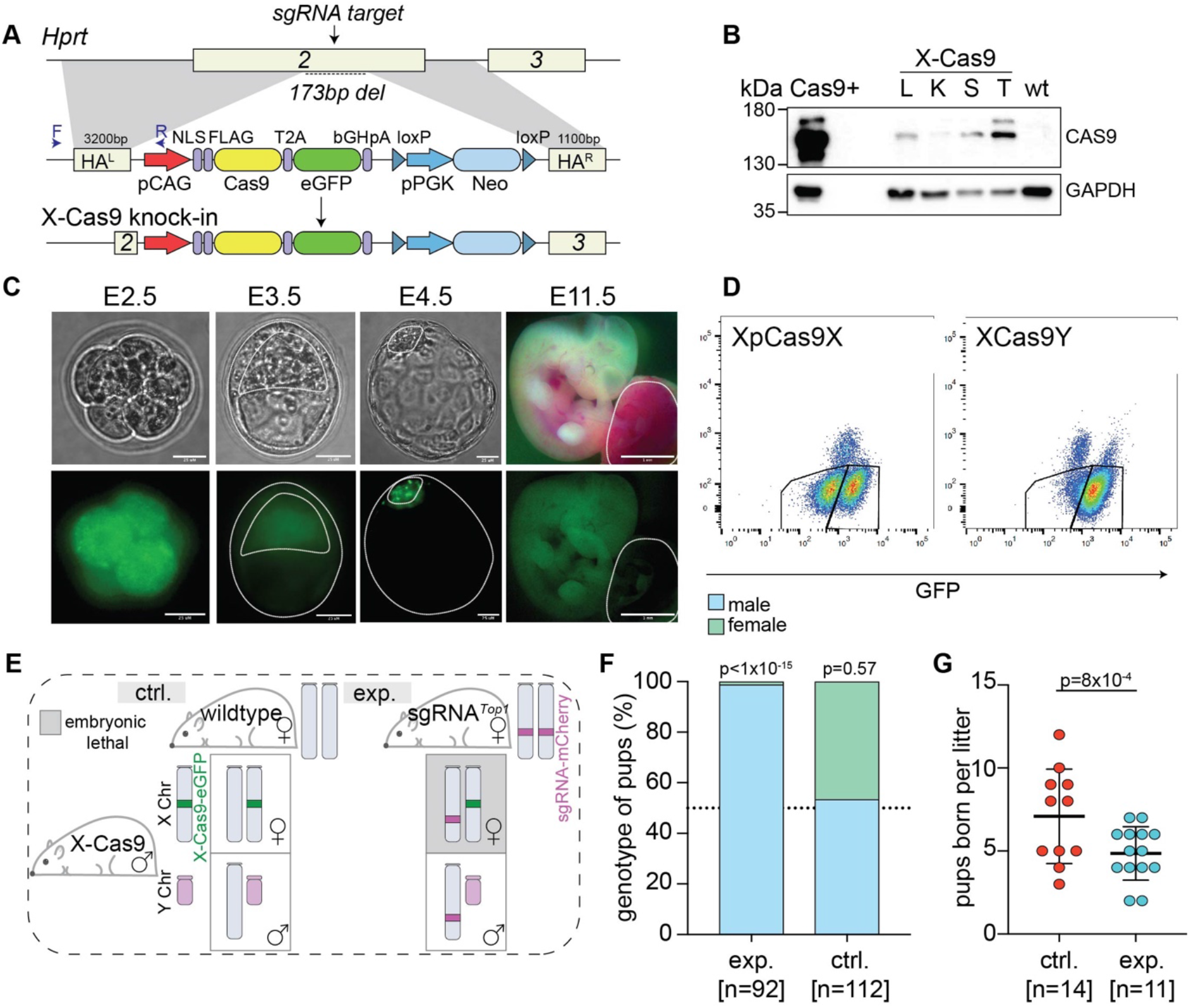
Generating the X-Cas9 transgenic line. (A) Schematic of X-Cas9 knock-in strategy. HA^L^; homology arm left, HA^R^; homology arm right, NLS; nuclear localisation signal, F; forward primer, R; reverse primer. (B) Western blot of X-Cas9Y tissues, wildtype and *R26*-Cas9 (Cas9+). L; liver, K; kidney, S; spleen, T; testis. Expected size: 158 kDa (CAS9) and 37 kDa (GAPDH). (C) eGFP expression in female XpCas9X embryos, with different lineages delineated by lines. E2.5 (n=22), E3.5 (n=4), E4.5 (n=3), sb; 25 μM. E11.5 (n=10) sb; 100 μM. (D) Flow cytometry from XpCas9X embryos (n=9) and XCas9Y embryos (n=2). (E) Schematic of the mating strategies, ctrl; control, exp; experimental. (F) Sex genotyping of pups born from control or experimental mating. Number of pups genotyped in brackets. P value: Chi-squared test, assuming 1:1 ratio of ‘expected’ offspring if co-inheritance of CRISPR-Cas9 alleles was not lethal. (G) Litter size. Number of litters in brackets. P value: Mann-Whitney test.

X-Cas9Y F2 transgenic males were viable and fertile, with testis weights comparable to wildtype males (102.3mg versus 104.2mg; 11 weeks old; p=0.67 Mann-Whitney test). We established by Southern blotting and digital droplet qPCR that the construct was present as a single copy (Supp Fig 2C;D). Expression of eGFP was observed in adult organs by fluorescence microscopy (Supp Fig 2E) and qPCR (Supp Fig 2F), and Cas9 expression was confirmed in adult organs by western blotting (Fig 3B).

Before assessing whether a paternally-inherited X-Cas9 (XpCas9) transgene caused lethality in daughters, we examined its expression in female (XpCas9X) pre-implantation embryos. In mice, the paternal X chromosome is initially active, before being silenced from the 4-8 cell stage by imprinted X-chromosome inactivation (XCI)^26,27^. Imprinted XCI is retained in the trophectoderm, but is reversed in the epiblast, after which random XCI ensues, giving rise to mosaic X-chromosome expression patterns^26,27^. Expression of eGFP recapitulated the known dynamics of paternal X expression, suggesting that the X-Cas9 transgene was subject to imprinted XCI. At E2.5 (8-16 cell stage) eGFP expression was observed (Fig 3C). At E3.5 (blastocyst stage), expression was reduced in the trophectoderm, where imprinted XCI is sustained, but was higher in cells of the inner cell mass, where X-chromosome reactivation takes place (Fig 3C). The reduction in trophectoderm eGFP expression was likely a result of imprinted XCI, because it was not observed in control female embryos carrying the transgene on the maternal X chromosome, which is not subject to XCI (Supp Fig 2G). During later development in XpCas9X embryos, silencing of the X-Cas9 transgene persisted in extraembryonic lineages, with eGFP undetectable in the trophectoderm at E4.5 and the placenta at E11.5 (Fig 3C). However, expression increased in the presumptive epiblast at E4.5, and persisted in the embryo proper at E11.5 (Fig 3C).

To determine if the X-Cas9 transgene was also subject to random XCI, we performed flow cytometry on cells derived from post-implantation XpCas9X female embryos, with XCas9Y male embryos as controls. If X-Cas9 was subject to random XCI, approximately half the XpCas9X cells should express eGFP, but if it escaped random XCI, all XpCas9X cells should express eGFP. In XpCas9X embryos, 47.3% of cells were eGFP-positive and 52.7% were eGFP-negative (n=9 embryos; Fig 3D). In male XCas9Y embryos (n=2), 86% of cells were eGFP positive (Fig 3D). The X-Cas9 transgene is therefore subject to random XCI.

To assess whether we could generate single-sex litters, X-Cas9Y hemizygous males were mated with either wildtype females (control mating) or homozygous sgRNA*^Top1^* females (experimental matings; Fig 3E). From control matings, male and female pups were recovered in approximately equal proportions (60M:52F; n=112 pups; non-significant deviation from Mendelian sex ratio, p=0.57; Fig 3F). In contrast, from the experimental mating, there was a striking sex skew, with 99% of pups being male (n=91/92, statistically significant deviation from Mendelian sex ratio, p<1×10^−15^, Fig 3F). Genotyping and low-pass whole genome sequencing revealed that the single, exceptional female was XO, a genotype that arises spontaneously in our mouse colony at a frequency of approximately 1:100 females. Intriguingly, this XO female had inherited a maternal X chromosome but no paternal X chromosome, and thus lacked the X-Cas9 transgene necessary to induce lethality (Supp Fig 2H). We conclude that co-inheritance of the X-Cas9 and sgRNA*^Top1^* induces lethality in all XX females.

Given the loss of female embryos, we predicted that the litter size in our experimental matings would be 50% of that in control matings. Intriguingly however, while the mean litter size was indeed reduced (4.6 versus 7.5, respectively, p=8×10^−4^; Fig 3G), it was 61% rather than 50% of controls. These findings, which were reminiscent of those observed in our *R26*-Cas9 and sgRNA*^Top1^* experimental matings (Fig 2J), again reveal the existence of a compensatory mechanism operating in the pre-implantation period to increase embryo number.

## Discussion

In this study we show that co-inheritance of a paternal X-linked Cas9 transgene and a maternal *Top1*-targeting sgRNA induces embryonic lethality in XX females, thereby generating male-only litters.

The CRISPR-Cas9 bicomponent system we describe here is superior to some pre-existing methodologies to generate single-sex litters. One of these methods is CRISPR-Cas9 gene drive. First trialled in mosquito models by targeting the female-specific *doublesex* splice variant, female offspring showed an intersex phenotype and were sterile, causing population collapse^28^. This study was expanded to a sex distorter gene drive technology, whereby a synthetic gene drive was designed to spread the X-shredding I-Ppol endonuclease at above-Mendelian frequency, resulting in male-only populations^29^. However, proof-of-principle CRISPR-Cas9 gene drives performed in the mouse remain largely inefficient^30^. We propose that our system has greater functionality for generating all-male litters in mammalian models.

Our proof-of-principle approach could be readily translated to the laboratory mouse model. The X-linked Cas9 and autosome-linked sgRNA*^Top1^* transgenic stocks can be maintained as mono-allelic lines and bred when necessary to generate single-sex litters. Conversely to gene drive CRISPR-Cas9 strategies, the risk of mutational resistance at the sgRNA-target occurring is not problematic, because the mono-transgenic lines are maintained independently and only combined when necessary. The production of single-sex litters using these transgenic lines for studies such as behaviour or reproductive science, will immediately reduce the production of the unrequired sex, transforming laboratory approaches to the 3Rs. The *Top1* sgRNA sequence is highly conserved between mice and many agricultural species and may therefore be adapted for use in the agricultural industry. Moreover, the method could be easily modified, with Cas9 integrated on the Y chromosome rather than the X chromosome. This alternative approach would permit the production of all-female litters. In our strategy, the sex-linked Cas9 transgene is lost in the embryonic-lethal population, producing surviving offspring that carry only the sgRNA. This approach may be preferable to that employed by Yosef et al^12^ in which the surviving sex carry and express a potentially harmful Cas9 endonuclease transgene.

Our data reveal that in mice a compensation mechanism operates during pre-implantation development that increases the number of pups by approximately one per litter. We speculate that this compensation is possible because there is an overproduction of fertilised zygotes compared with uterine capacity^31^. Thus, loss of embryos prior to implantation could be buffered by implantation of embryos from this excess pool. This reveals an unexpected benefit of our targeting system that could increase the number of the desired sex over that derived from control matings

Finally, our CRISPR-Cas9 bicomponent system could be applied to other scenarios in which mutations are required in a sex-specific manner. Many harmful mutations, e.g. those causing cancer, are assayed preferentially in one biologically-relevant sex^32,33^, yet the unrequired sex also suffers the ill effects of this mutation. Our technology would reduce suffering in the unrequired sex, in line with the 3Rs.

## Methods

### Maintenance of mouse lines

All mouse lines were maintained with appropriate care according to the United Kingdom Animal Scientific Procedures Act (1986), UK Home Office, and the ethics guidelines of the Francis Crick Institute. All mouse lines used were strain *Mus musculus*. All wildtype mice used were C57BL/6J. X-Cas9 transgenic mice were generated on a C57BL/6N mESC background, and then maintained on a C57BL/6J background, after generating a stable, germline transmitting line. The *H11*-attPX3 mice were backcrossed to at least seven generations of C57BL/6J by Charles River, prior to purchase for zygotic microinjection. The *H11*-sgRNA*^Top1^* mouse line was also maintained on a C57BL/6J background. Litter mate controls were used where possible. All mice were kept in IVC cages, with constant access to food, automatic watering systems, and air management systems which control air flow, temperature and humidity. The mouse lines were checked on a daily basis, and were maintained in specific pathogen free (SPF) conditions.

### Embryonic stem cell derivation and maintenance

All mESC lines were maintained in 2i/LIF conditions on laminin-coated tissue culture grade plasticware^34^. To derive mESCs, embryos were collected at E3.5 by flushing the uterus with Follicle Holding Medium (FHM) from timed mating 6-8 week old females. Embryos were placed in individual wells of a 24-well plate with 500ul of 2i/LIF. Outgrowths were dissociated and mESCs seeded into a 4-well plate in 2i/LIF. mESCs were passaged by removing 2i/LIF, washing with PBS, followed by trypsinisation with TrypLE (Gibco), quenching with 2i/LIF and pipetting into a single cell suspension. Following centrifugation at 200 g for 3 mins, mESCs were resuspended and seeded in new plates^35^.

### Fluorescence activated cell sorting (FACS)

Transfected mESCs were trypsinised using TrypLE into a single cell suspension, centrifuged at 200 g for 3 mins, and resuspended in sorting media (2% FBS in 2i/LIF). mESCS were filtered (40 μM) and sorted using the Aria Fusion Flow Cytometer with a 100 μM nozzle. mESCS were firstly gated on forward and side scatter properties, followed by gating on either eGFP+ single-positive only or eGFP+mCherry+ double positive expression. The eGFP-only population acted as the CRISPR-Cas9 negative control.

### Embryo dissociation and flow cytometry

E11.5-E12.5 embryos were dissociated and prepared for flow cytometry according to previously published protocols^36^. Dissociated cells were filtered (40 μM) and maintained on ice in sterile PBS with 2% FBS prior to flow cytometry and analysis on the MACSQuant VYB. Single cells were analysed on forward and side scatter properties, followed by gating on GFP expression.

### sgRNA design

All sgRNAs were designed using publicly available *in silico* tools^37^. Single sgRNAs with a predicted high on-target activity and low off-target activity were selected. Oligonucleotides with BbsI overhangs were annealed and ligated into the relevant vector, according to published protocols^38^.

### Generating the pLethal/TARGATT mouse line

The pLethal targeting vector was generated using pX333 (addgene #64073)^39^; replacing the Cas9 cassette with an mCherry reporter. Individual sgRNAs were cloned into pLethal using BbsI^38^. The knock-in targeting vector was generated by cloning the pLethal U6-sgRNA cassettes and pCbh-mCherry reporter into the TARGATT MCS vector #3^19,20^ (Applied StemCell). The TARGATT vector was microinjected into attPx3 embryo pronuclei with ϕC31 integrase, and embryos were surgically transferred into pseudopregnant females. Founders were screened by *in vivo* fluorescence imaging at 3-4 days post birth using the IVIS Lumina XR (Caliper LifeSciences) with “Living Image 4.4” software, excitation filter at 535nm and emission filter dsRed.

### Generating the X-Cas9 mouse line

X-Cas9 targeting vectors were generated using the pX330 (addgene #42230)^40^ plasmid backbone, containing a pCAG driven 3X FLAG-NLS-Cas9-T2A-eGFP construct. X chromosome homology arms, amplified from C57Bl/6J DNA, and a LoxP-flanked pPGK-Neomycin cassette were inserted using directional cloning or Gibson Assembly (NEBuilder HiFi DNA Assembly Cloning Kit). C57BL/6N mESCs were maintained in serum/LIF conditions and transfected with X-Cas9 targeting vector plasmid and an sgRNA targeting *Hprt* exon 2 using lipofectamine 2000, according to manufacturer’s instructions. Targeted mESC clones were selected by G418 (270 mg/ml) for 8-10 days. Surviving clones were picked into a 96-well plate and expanded. Expanded mESC lines were lysed by the addition of Bradley Lysis buffer (1 M Tris-HCl, 0.5 M EDTA, 10% SDS, 5M NaCl) and proteinase K (1 mg/ml) digestion. DNA was precipitated by the addition of ice cold EtOH/NaCL (100% EtOH, 5M NaCl). PCR genotyping was performed on extracted DNA in a total volume of 25 μl (12.5 μl NEB Q5 High-Fidelity Master Mix, 10 mM each primer), utilising primer forward and reverse pairs aligning to the endogenous *Hprt* locus and to the transgene construct. Resultant PCR amplicons were analysed by gel electrophoresis for corresponding to the expected amplicon size, and by Sanger sequencing. Targeted mESC clones were injected into albino C57BL/6J blastocysts, surgically transferred into pseudopregnant females, and left to litter. X-Cas9 mESC contribution to founders was assessed by coat colour. High contribution transgenic males were bred with C57BL/6 albino females, and offspring with black coat colour were genotyped for the transgene, to confirm germline transmission.

### Genotyping offspring from breeding CRISPR-Cas9 stocks

Pups born from the control and/or experimental breeding programmes were genotyped using assays for the Cas9-eGFP transgene, the mCherry transgene, and sexed by the presence of Y-linked gene *Sry*. X-Cas9 hemizygous versus homozygous females were distinguished by genotyping for the *Hprt* exon 2 deletion. The XmO female generated from breeding X-Cas9 males with *H11*-sgRNA*^Top1^* homozygous females, was characterised by DNA extraction from ear biopsy tissue; X-chromosome and transgene copy number analyses, and low-pass whole genome Nanopore sequencing.

### Primer design

All primer pairs used in this study were designed using the publicly available tool Primer3 (http://bioinfo.ut.ee/primer3/). To amplify the target *Top1* exons for sequencing, oligonucleotide forward and reverse primers were edited to contain MiSeq adaptor sequences.

### DNA extraction, MiSeq high throughput sequencing and indel analysis

Samples, e.g. mESCs, embryos or tissue, were lysed by the addition of lysis buffer (10X KT buffer, 10% NP40) with proteinase K (1 mg/ml) digestion. Correct amplification of *Top1* exons was confirmed by gel electrophoresis. The PCR amplicons were purified using solid-phase reversible immobilisation beads^41^, and underwent library preparation (Illumina Nextera Index Kit V2), following by a second purification using Agencourt AMPure beads. The purified DNA library was quantified, normalised and pooled prior to sequencing on the Illumina MiSeq platform to generate paired-end (2 x 250 bp) sequencing reads. Resultant reads were demultiplexed and fastq files were collapsed using FastX Toolkit (v0.0.13; https://github.com/agordon/fastx_toolkit). To assess the rate of indel-production by CRISPR-Cas9, the reads were aligned to the mouse reference genome mm10 with the Burrows-Wheeler Alignment tool (BWA, v0.7.170)^42^ using the *mem* algorithm with default settings and then analysed using the R package CrispRVariants (v1.14.0)^43^. Scripts are deposited on github (https://github.com/jzohren/crispr-miseq).

### Low-pass whole genome sequencing and low-pass whole genome Nanopore sequencing

DNA was extracted by the phenol-chloroform method, as described previously^44^. Samples underwent library preparation using the Illumina Nextera Flex protocol, according to manufacturer’s instructions. Libraries were sequenced to achieve approximately 0.1X coverage per sample. Low-pass whole genome sequencing reads were aligned to *mm10* using BWA, with the number of reads mapped extracted from the data. For Nanopore sequencing, DNA was extracted by the phenol-chloroform method. Samples were prepared according to the Oxford Nanopore Technologies (ONT) SQK-LSK109 library preparation protocol. Libraries were sequenced on a FLO-MIN106D flow cell on the MinION. Basecalling was performed using ONT-Guppy v3.2, and data was mapped using minimap2^45^ and SAMtools^46^.

### Quantitative PCR analysis

RNA was extracted using TRI Reagent (Sigma-Aldrich), according to manufacturer’s protocol. cDNA was synthesised using the Thermo Scientific First Strand cDNA Synthesis Kit, according to manufacturer’s protocol. Samples were analysed in triplicate, in 10 μl total volume (5 μl TaqMan 2X Universal PCR Master Mix, 0.5 μl TaqMan probe, 2.5 μl nuclease-free water, 2μl cDNA). Resulting ddCt values were calculated by normalising to *Gapdh* expression from C57BL/6 samples.

### Digital Droplet qPCR

DNA was extracted by phenol-chloroform precipitation. Digital droplet qPCR (ddPCR) reactions were performed in 20 μl total volume with 20 ng DNA, according to manufacturer’s instructions (Bio Rad ddPCR Supermix for Probes). The ddPCR was performed in a Bio Rad PCR machine, and analysed using QuantSoft.

### Protein extraction and western blot

Protein was extracted from samples using 1X RIPA buffer with additional phosphatase and protease inhibitors, and PMSF. Upon adding protein extraction buffer to samples, samples were kept on ice for 30 minutes, following centrifugation at 8,000 rpm at 4 °C for 10 minutes. Supernatant was collected and protein quantified using a bicinchoninic acid (BCA) assay and analysed using Kaleido 2.0. Proteins were separated using PAGE system and transferred to 0.45 μm pore Nitrocellulose membrane (Amersham Protran). Membranes were blocked with 5% skimmed milk/TBST for 1h at room temperature and incubated with primary antibodies overnight at 4°C. CAS9 and TOP1 antibodies were used at 1:500, α-Tubulin at 1:2000, GAPDH at 1:3000 dilutions. Appropriate secondary antibodies conjugated to HRP were used and signals were detected using Clarity Western ECL Substrate (Bio-Rad).

### Southern blot

DNA was extracted by phenol-chloroform precipitation, digested using appropriate restriction enzymes, and phenol-chloroform precipitation repeated. DNA was loaded onto a 1% agarose gel and gel electrophoresis run overnight at 29V, followed by addition of bromophenol blue, and further running at 50V for 2-3 hours. Following gel electrophoresis, the agarose gel was treated by washing in depurination (0.25M HCl), denaturation (1.5M NaCl, 0.5M NaOH) and neutralisation (1.5M NaCl, 0.5M Tris pH 7.5) buffers and overnight blotting onto a positively-charged nylon membrane. After blotting, the DNA was fixed by UV crosslinking (1200U joules, 2 minutes) and drying. The membrane then underwent hybridisation to the Neomycin probe, produced according to manufacturer’s instructions (Roche DIG probe synthesis kit) and incubation overnight in a hybridisation oven at the optimal temperature (48 °C for Neomycin). Post-hybridisation, the membrane was washed (2X SSC, 0.1% SDS) at room temperature, and at 65 °C (0.1X SSC, 0.1% SDS). Following this, the membrane was blocked with blocking buffer and incubated with anti-DIG antibody (Roche detection kit), washed (maleic acid, 0.3% tween-20), and exposed to CSPD in detection buffer under darkness before film development.

## Acknowledgements

The authors thank the Francis Crick Institute Genetic Modification Service (GeMS), Flow Cytometry, Thomas Snoeks (*In Vivo* Imaging), Biological Research and Advanced Sequencing facilities for their expertise; Tatyana Nesterova and Neil Brockdorff (University of Oxford) for stem cell targeting advice, and members of the J.M.A.T lab for comments and discussion on the manuscript.

## Author Contributions

J.M.A.T and P.J.I.E conceived the project. J.M.A.T and C.D. designed the project. C.D. performed the molecular biology, Southern blotting, embryonic stem cell experiments, embryo experiments, fluorescence imaging, mouse colony genotyping and phenotyping, and wrote the manuscript. V.M. performed the western blotting. J.Z. performed bioinformatic analysis. D.M.S performed the low-pass Nanopore whole genome sequencing, and provided advice on experimental design. O.A.O. managed mouse colonies and performed genotyping.

