## Supplemental Figures for "Generating single-sex litters: development of CRISPR-Cas9 genetic tools to produce all-male offspring"

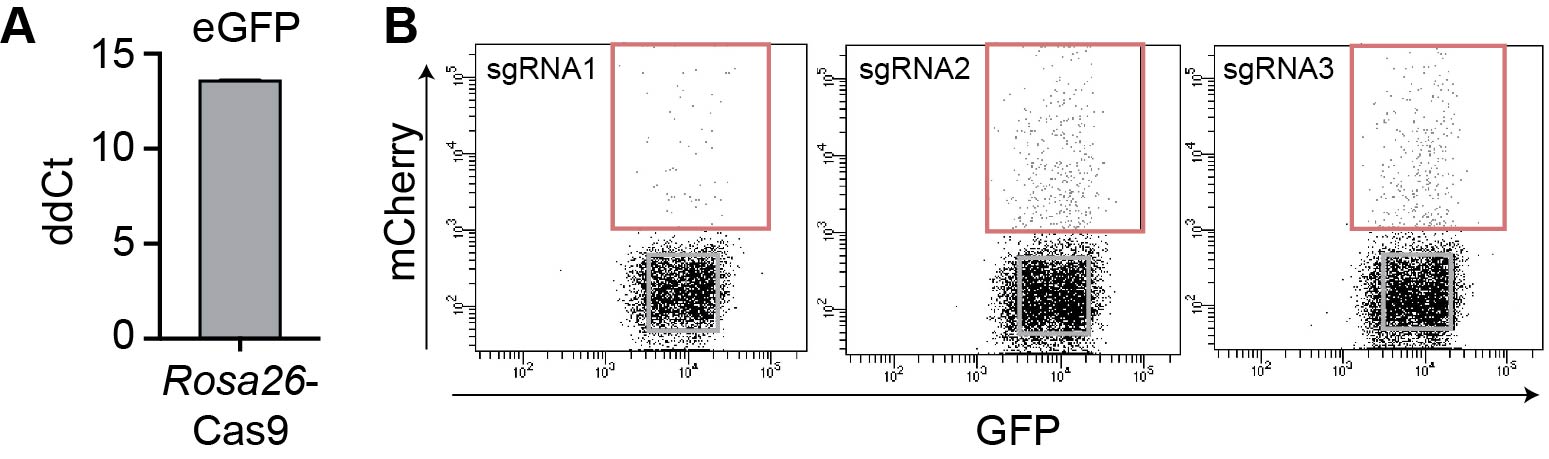

**Supp Fig 1.**

(A) Quantitative PCR of eGFP expression in *R26*-Cas9 mESCs, normalised to *Gapdh* in a wildtype sample. (B) Fluorescence activated cell sorting of eGFP/mCherry double-positive mESCs (red) and eGFP+ single-positive mESCS (grey).

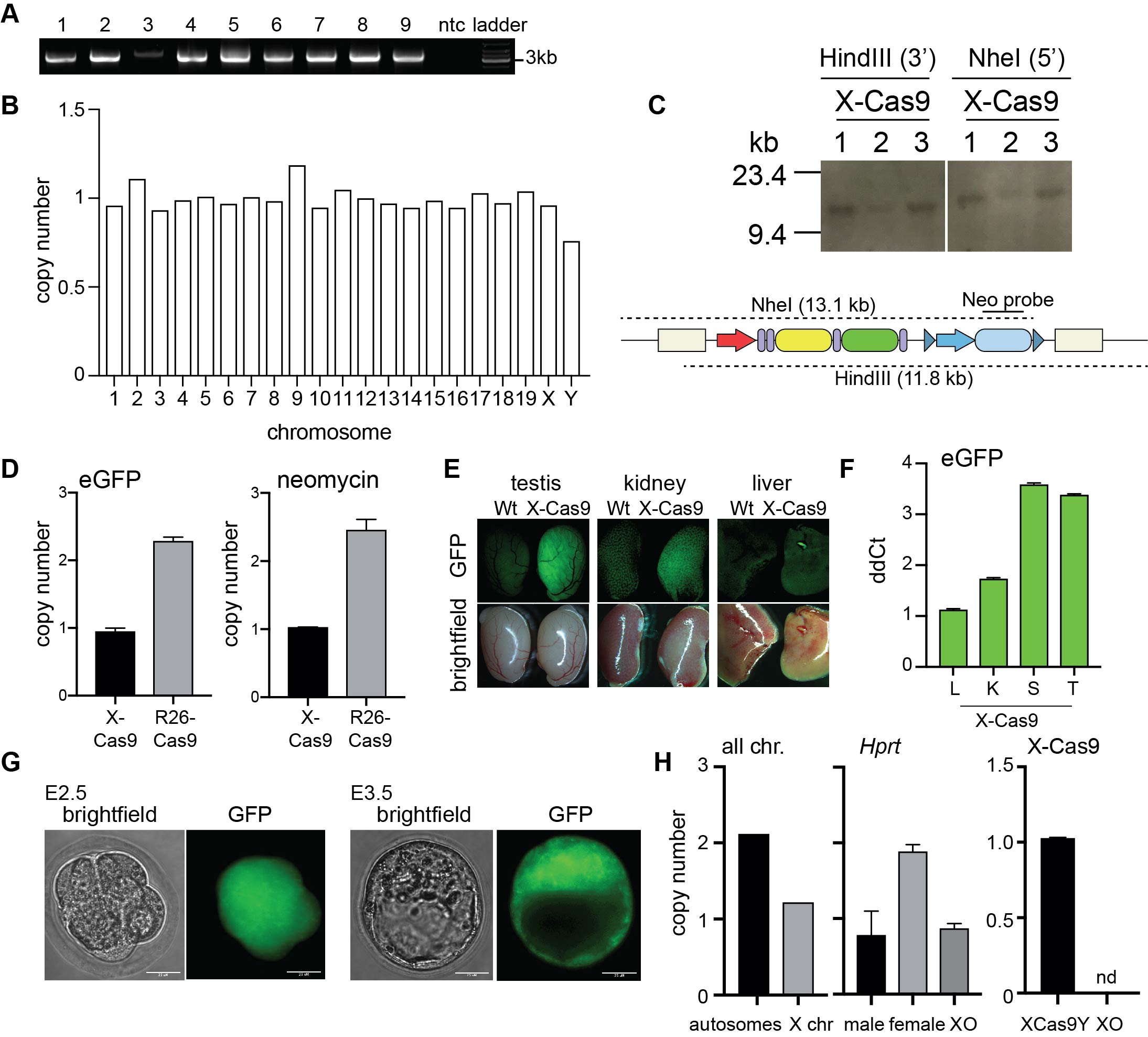

**Supp Fig 2.**

(A) PCR genotyping 5’ boundary PCR. ntc; no template control. Expected size: 3.5 Kb.(B) Low-pass whole genome sequencing of X-Cas9 clone 5 mESCs. Normalised to C57Bl/6J XY. (C) Southern blot. Expected size: HindIII 11.8kb, NheI 13.1kb. (D) Digital droplet PCR of eGFP and Neomycin. Normalised to *Tfrc.* Error bars: s.d. (n=3). (E) Fluorescence microscopy of X-Cas9Y tissues. (F) Quantitative PCR of eGFP expression in X-Cas9Y, normalised to *Gapdh* in wildtype sample. L; liver, K; kidney, S; spleen, T; testis. (G) Fluorescence microscopy of XmCas9X embryos. Scale bar: 25 μM. (H) Low-pass whole genome sequencing (all chromosomes) and ddPCR (*Hprt,* X-Cas9). ddPCR samples normalised to *Tfrc*. nd; not detected. Error bars: s.d. (n=3).
